# Intravital calcium imaging of meningeal macrophages reveals niche-specific dynamics and aberrant responses to brain hyperexcitability

**DOI:** 10.1101/2025.10.01.679335

**Authors:** Simone Carneiro-Nascimento, Chao Wei, Anna Gutterman, Dan Levy

**Affiliations:** Department of Anesthesia, Critical Care and Pain Medicine, Beth Israel Deaconess Medical Center, Harvard Medical School, Boston, United States

**Keywords:** Meningeal macrophages, calcium imaging, dural vasomotion, cortical spreading depolarization, migraine

## Abstract

The meninges, which envelop and protect the brain, host a dense network of resident macrophages with diverse roles in regulating homeostasis and neuroinflammation. Despite their importance, we have a limited understanding of their behavior in vivo. Many dynamic cellular functions of macrophages involve intracellular Ca^2+^ signaling. However, virtually nothing is known about the spatiotemporal Ca^2+^ dynamics of meningeal macrophages in vivo. We developed a chronic intravital two-photon imaging approach and related computational analysis tools to interrogate meningeal macrophage Ca^2+^ dynamics, at subcellular resolution, in a novel Pf4^Cre^:TIGRE2.0^GCaMP6s/wt^ reporter mouse model. Using imaging in awake mice, we characterized Ca^2+^ activity in meningeal macrophages at steady state and in response to cortical spreading depolarization (CSD), an aberrant pro-inflammatory brain hyperexcitability event implicated in migraine, traumatic brain injury, and stroke. In homeostatic meninges, macrophages in the dural perivascular niche exhibited several Ca^2+^ dynamic features, including event duration and signal frequency spectrum, distinct from those of localized to the interstitial, non-perivascular niche. Simultaneous tracking of macrophage Ca^2+^ dynamics and local vasomotion revealed a subset of dural perivascular macrophages whose activity was coupled to locomotion-driven diameter fluctuations of their associated vessels. Most perivascular and non-perivascular meningeal macrophages displayed propagating intracellular Ca^2+^ activity and synchronized intercellular Ca^2+^ elevations, potentially driven by extrinsic factors. In response to CSD, the majority of perivascular and non-perivascular meningeal macrophages showed a persistent decrease in Ca^2+^ activity, while a smaller subset displayed Ca^2+^ elevations. Mechanistically, CGRP/RAMP1 signaling mediated the increase but not the decrease in CSD-mediated Ca^2+^ signaling. Collectively, our results highlight a previously unknown diversity of Ca^2+^ dynamics in meningeal macrophages at steady state and in response to an aberrant brain hyperexcitability event linked to neuroinflammation.

## Introduction

Resident macrophages are key myeloid immunocytes that play an important role in innate immune surveillance and defense across various peripheral tissues and organs (*1–3*). The central nervous system also harbors a large subset of parenchymal macrophages, known as microglia, and several distinct subsets of macrophages localized to the brain’s border tissues, including the choroid plexus, perivascular spaces, and the meningeal compartments that cover, protect, and support the brain (*4–8*). Macrophages are the predominant immune cell type within the brain meninges, and recent studies have demonstrated their diverse ontogeny, transcriptomic profiles, and immune functions at steady state (*5, 8–12*) and in several neuropathological conditions (*8, 13–16*).

Cytoplasmic calcium (Ca^2+^) signaling underlies a wide variety of cellular homeostatic and inflammatory processes in macrophages (*17–23*). In addition to intracellular Ca^2+^ elevation, distinct spatiotemporal dynamics - including oscillation patterns, intracellular propagations, and intercellular synchronization of Ca^2+^ signals - may regulate different macrophage functions during steady state and pathophysiology (*20–25*). Despite our increased understanding of the diverse molecular signatures and contributions of meningeal macrophages to homeostasis and neuroinflammation, virtually nothing is known about their Ca^2+^ signaling heterogeneity in both healthy and diseased states.

Here, we comprehensively characterized the Ca^2+^ dynamics of individual macrophages localized to the brain meninges by combining intravital two-photon Ca^2+^ imaging in a novel reporter mouse line, in which the Ca^2+^ reporter GCaMP6s is expressed in platelet factor 4 (Pf4^+^) meningeal macrophages, with an event-based signaling analysis pipeline. Our data reveal several distinct spatiotemporal Ca^2+^ dynamic features in perivascular versus interstitial non-perivascular meningeal macrophages, including a unique coupling between the Ca^2+^ signals of dural perivascular macrophages and behaviorally-driven vasomotion of their associated dural vessels at steady state. Furthermore, our data uncover both increases and decreases in Ca^2+^ activity in distinct subsets of meningeal macrophages in response to cortical spreading depolarization (CSD), a pathophysiological brain hyperexcitability event linked to headache pain and neuroinflammation (*26*) in migraine, traumatic brain injury, and stroke (*27*). Mechanistically, our data suggest that calcitonin gene-related peptide/receptor activity-modifying protein 1 (CGRP/RAMP1) axis mediates CSD-evoked macrophage Ca^2+^ elevation and related brain-to-meninges neuroimmune signaling pathway, potentially involving CGRP released from sensitized meningeal sensory neurons acting on macrophage CGRP receptors (*16, 28, 29*).

## Results

### Characterizing macrophage Ca^2+^ signaling features in homeostatic brain meninges

Previous intravital imaging studies exploring the spatiotemporal dynamics of tissue-resident macrophages have primarily used CX3C motif chemokine receptor 1 (CX3CR1)-based mouse reporter strains (*13, 22, 30, 31*). However, other resident monocyte-derived cells are labeled in these reporter mice (*32, 33*). Moreover, brain microglia also express CX3CR1 (*13, 33–35*), limiting the use of these reporter mice for resolving the spatiotemporal subcellular Ca^2+^ dynamics of macrophages localized to the relatively thin meningeal layers covering the brain parenchyma. To systematically characterize meningeal macrophage Ca^2+^ dynamics *in vivo* and avoid contamination from Ca^2+^ signals arising from superficial parenchymal microglia, we leveraged recent findings showing that Pf4 is highly enriched in meningeal macrophages but not in other meningeal immunocytes or parenchymal microglia (*10, 16, 36*), and generated transgenic reporter mice expressing the highly sensitive Ca^2+^ indicator GCaMP6s (using TIGRE2.0-based Ai162D mice (*37*)) in Pf4^+^ macrophages (using Pf4^Cre^ mice (*38*)). Notably, 100% of meningeal macrophages are labelled in Pf4^Cre^-based reporter mice (*36*).

Anesthetic agents impact intracellular Ca^2+^ signaling, including in brain macrophages and other non-excitable glial cells (*39–41*). We therefore investigated subcellular Ca^2+^ activity of meningeal macrophages in awake, behaving mice. We imaged meningeal macrophages via a chronic cranial window implanted together with a restraining headpost over the intact dura mater overlying the posterior neocortex. This chronic window approach produces minimal inflammatory responses in the cortex and meninges below the window (*42, 43*). After at least 7 days of recovery, mice were gradually habituated across multiple days to head restraint while free to run on a wheel (**Figure 1b**). We tracked Ca^2+^ transients of GCaMP6s-labeled meningeal macrophages using high-speed two-photon microscopy in 37 fields of view (FOVs) from 7 mice. We first corrected the imaging movies for locomotion-evoked meningeal translational shifts using rigid registration (*43*). Movies were then processed using the AQuA2 data analysis platform (*44*), which implements an unbiased event-based approach to capture spatiotemporal Ca^2+^ event dynamics (**Figure 1C**). Based on spatial analysis, we assigned events (n=1361) with their corresponding Ca^2+^ features to each macrophage, using data from 503 cells.

**Figure 1.**
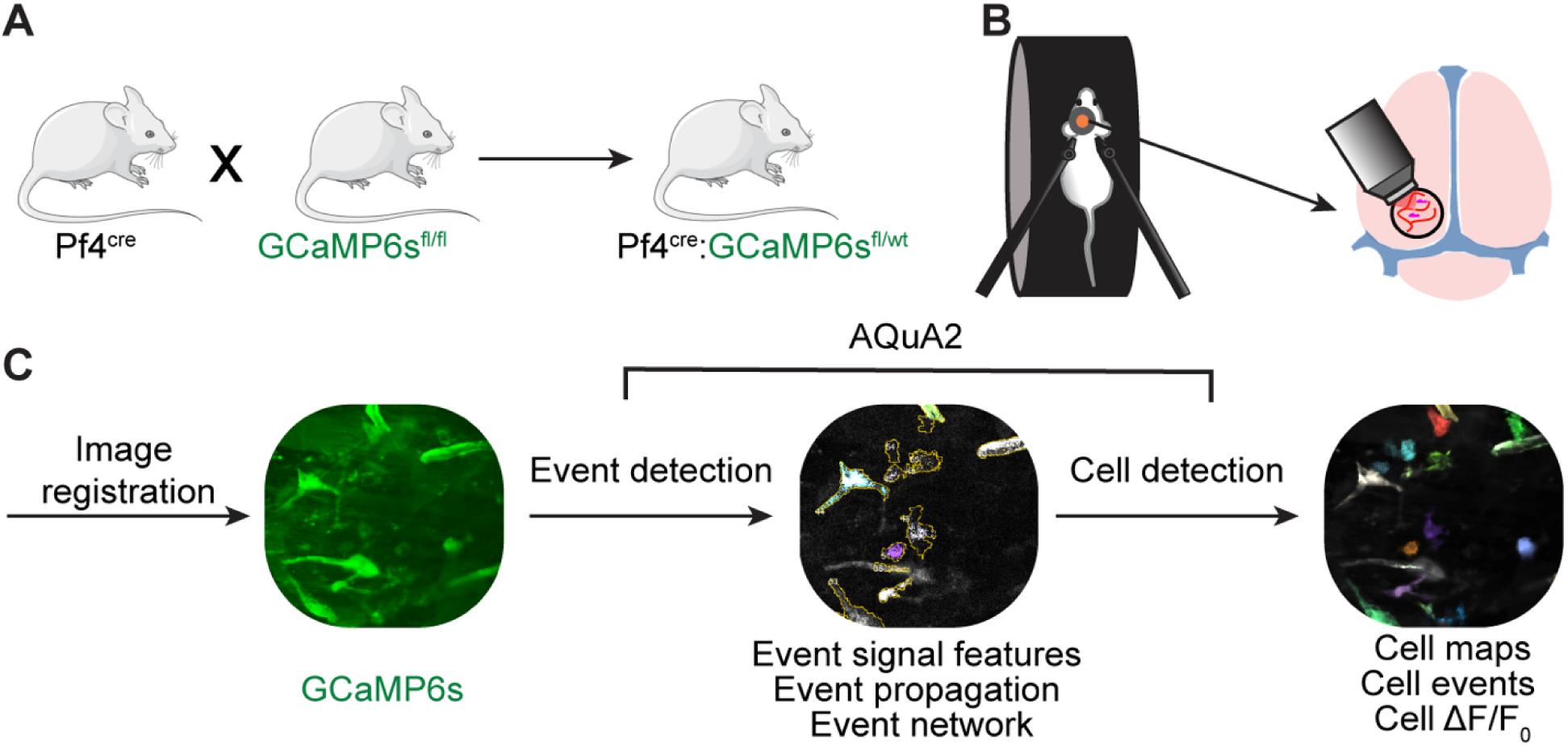
Imaging meningeal macrophage Ca^2+^ dynamics in awake behaving mice. **(A)** Pf4^Cre^:GCaMP6s^fl/wt^ reporter mouse construct for imaging meningeal macrophages Ca^2+^ activity.**(B)** Experimental procedure for two-photon imaging of meningeal macrophage Ca^2+^ activity. Following the implantation of a headpost and a cranial window, mice were habituated to head restraint and subjected to two-photon microscopy while head-fixed on a running wheel to study meningeal macrophage Ca^2+^ activity. **(C)** Macrophage Ca^2+^ imaging processing pipeline.

Meningeal macrophages occupy two distinct niches: perivascular (along the abluminal surface, physically contacting mural cells) and non-perivascular (within the interstitial space) (*8, 45*). These spatial distributions may dictate their divergent roles in meningeal immunity and vascular regulation. We therefore characterized the Ca^2+^ dynamics of these two anatomically distinct macrophage populations (perivascular, n=122; interstitial, non-perivascular, n=381, respectively, **Figure 2A and Video S1**). Most perivascular macrophages (93.4%, n=114) displaying ongoing Ca^2+^ activity were associated with vessels in the dura mater (labeled with a TRITC-Dextran tracer, see methods). The two meningeal macrophage subpopulations exhibited several distinct Ca^2+^ activity features. While the total area of Ca^2+^ activity in the peri- and non-perivascular macrophages was similar (**Figure 2B**), the signal perimeter of perivascular macrophages was significantly greater (**Figure 2C**) and exhibited a more elongated shape (**Figure 2A, 2D**), in agreement with their rod-shaped morphology (*8, 46*). The Ca^2+^ event duration in the perivascular macrophages was also longer compared to the interstitial macrophage subpopulation (**Figure 2F**). The peak Ca^2+^ activity level (Max ΔF/F_0_, **Figure 2E**) and event rate (**Figure 2G**) were, nonetheless, similar in the two meningeal macrophage subpopulations.

**Figure 2.**
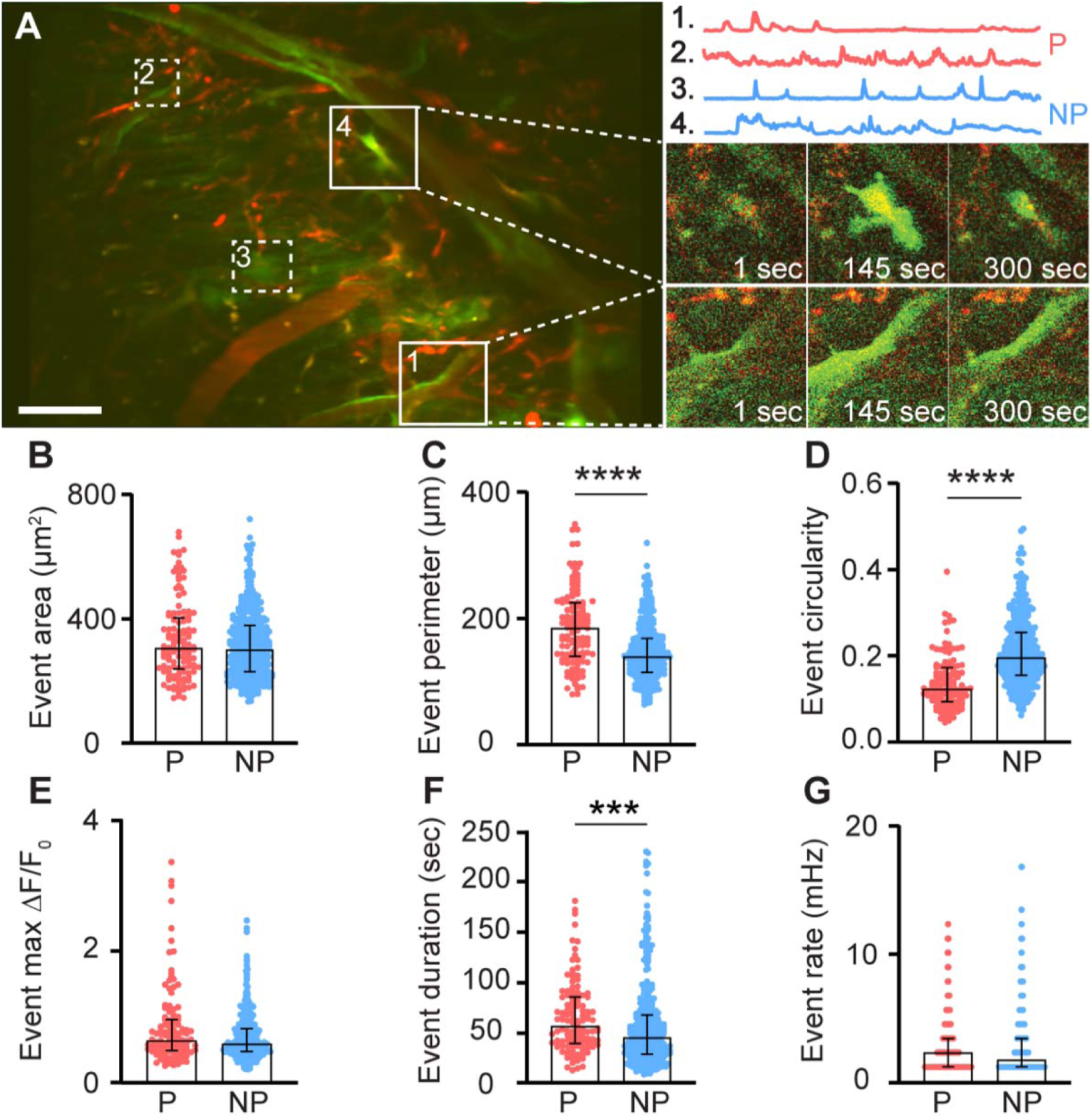
Ca^2+^ dynamic features of meningeal macrophages at steady state. **(A)** Left: Mean projection of an example FOV depicting perivascular (P, red; 1, 2) and non-perivascular (NP, blue; 3, 4) meningeal macrophages (white squares). Scale bar 50 μm. Right: Corresponding macrophages with representative 900-second Ca^2+^ activity traces (top) and their fluorescence signal at selected time points (bottom). **(B-G)** AQuA2-based morphological and Ca^2+^ event functional features of perivascular (P=122 cells) and non-perivascular (NP, n=381 cells) meningeal macrophage, n=37 fields of view (FOVs) from 7 mice. (B) Event area, (C) Event perimeter, (D) Event circularity, (E) Event max DF/F_0_, (F) Event duration, (G) Event rate. Data (B-G) represent median ± IQR. ***p<0.001, ****p<0.0001, Mann-Whitney U-test.

Distinct Ca^2+^ signal frequency spectra may underlie different biological functions of macrophages (*22*). We therefore analyzed meningeal macrophage Ca^2+^ signal waveforms using a multi-step signal processing and clustering analysis. We clustered cells based on two distinct patterns of Ca^2+^ activity. Cells in Cluster 1 (n=40) exhibited a more noisy-like activity pattern characterized by multiple frequencies, while cells in Cluster 2 (n=463) displayed a single dominant frequency at 0.01 Hz (**Figure 3A**). To further explore these cellular signaling differences, we combined the clustering data with AQuA2-derived features and observed that macrophages in Cluster 1 showed a larger event perimeter, but lower circularity and peak magnitude than those in Cluster 2 (**Figure 3D-F**). While Cluster 1 cells had a lower signal-to-noise ratio compared to Cluster 2 cells (**Figure 3I**), both Clusters displayed similar event area, duration, and rate (**Figure 3C, G, H**). Finally, we observed a significant association between cell cluster (1 vs. 2) and cell type, with Cluster 1 predominantly comprising perivascular macrophages and Cluster 2 comprising primarily non-perivascular macrophages (**Figure 3J**), further suggesting that these two meningeal macrophage subpopulations have distinct Ca^2+^ signaling properties.

**Figure 3.**
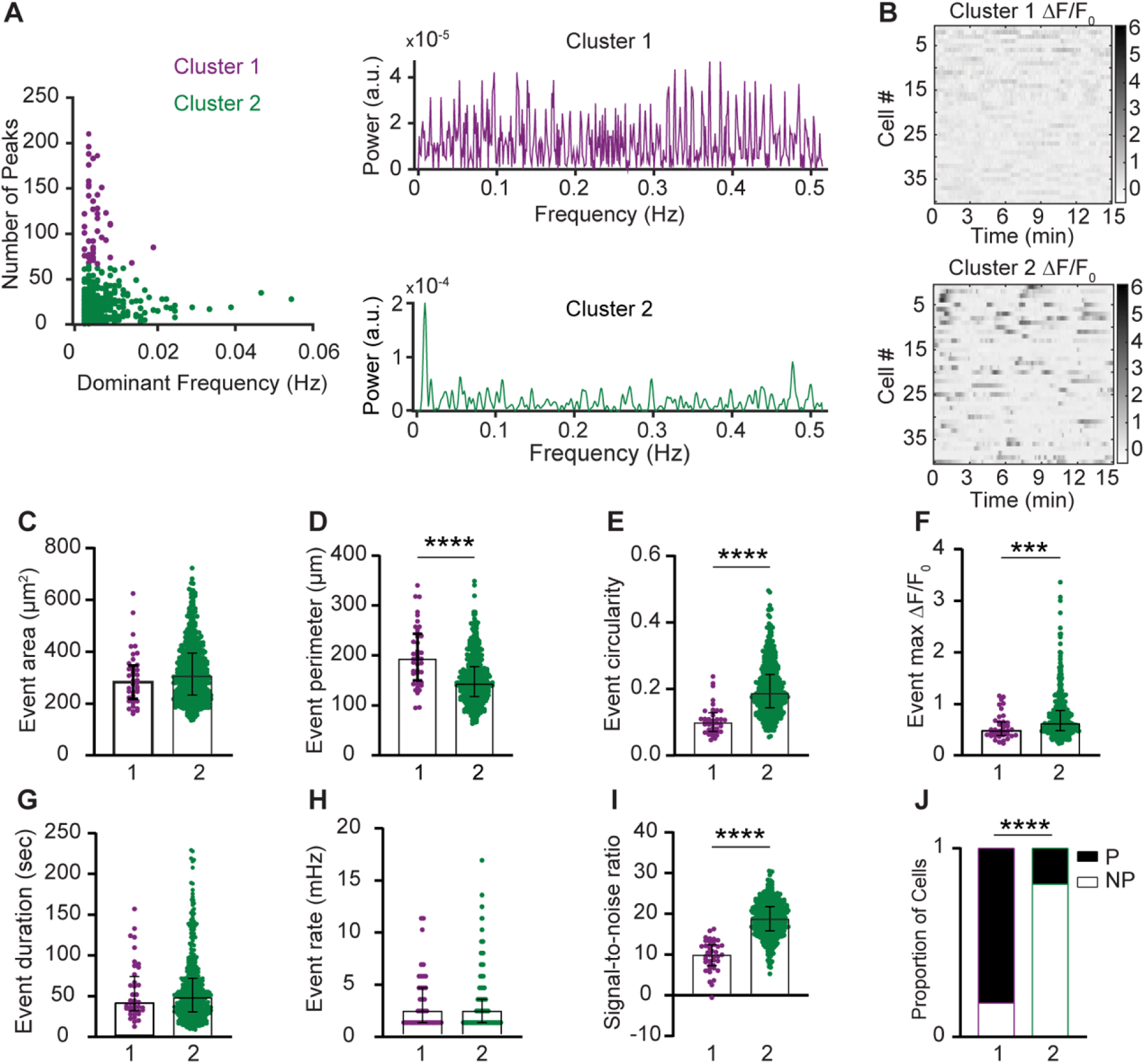
Intracellular Ca^2+^ signal frequency spectra of macrophage subsets in the steady-state meninges. **(A)** Left: Clustering of meningeal macrophage Ca^2+^ activity based on frequency-domain features and peak detection. Cluster 1 (purple, n=40 cells) and Cluster 2 (green, n=463 cells), n=37 fields of view (FOVs) from 7 mice. Right: Power spectrum density (PSD) of Ca^2+^ signals for each cluster. **(B)** Example DF/F_0_ heatmaps of Cluster 1 and Cluster 2 macrophages. **(C-H)** Morphological and Ca^2+^ functional features of Cluster 1 and Cluster 2 macrophages. (C) Event area. (D) Event perimeter (E) Event circularity. (F) Event max DF/F_0_. (G) Event duration. (H) Event rate. (I) Signal-to-noise ratio from Clusters 1 and 2. (J) Distribution of cell types across clusters. Data (C-I) represents median ± IQR. ***p<0.001, ****p < 0.0001, Mann-Whitney U-test Data (J) represents the cell proportion. ****p<0.0001, Chi-square test.

Intracellular Ca^2+^ signal propagation underlies diverse cellular functions and has been recently identified in macrophages *in vitro* (*21*) and in skin-resident macrophages *in vivo* (*47*). By assessing the Ca^2+^ signal propagation maps for each event within a defined cell, we identified two distinct patterns of activity: propagating events, in which Ca^2+^ signals traveled throughout the entire cell, and stationary events (**Figure 4A**). Propagating events showed varied signal source regions and directionality. Most macrophages (perivascular, 94.3%, n=115; non-perivascular, 86.9%, n=331) exhibited only propagating events, while a small minority of cells displayed a mix of propagating and stationary events or exclusively stationary activity events (**Figure 4B**).

**Figure 4.**
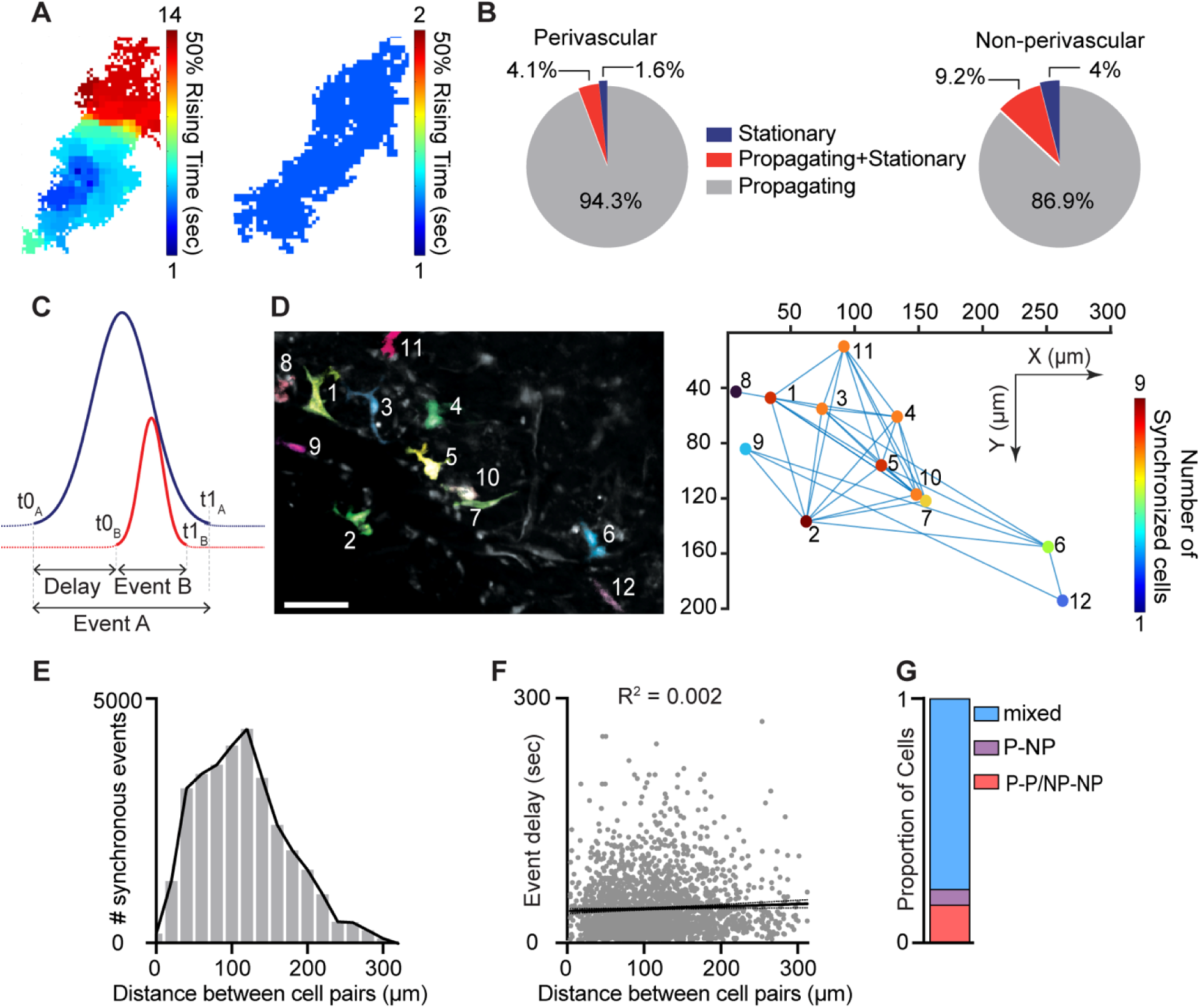
Intracellular propagation and intercellular synchronization of meningeal macrophage Ca^2+^ activity. **(A)** Spatial maps of two distinct Ca^2+^ events. Left: Propagating Ca^2+^ activity. Right: Stationary Ca^2+^ activity. **(B)** Distribution of event propagation profiles in perivascular (n=122 cells) and non-perivascular (n=381 cells) meningeal macrophages, n=37 fields of view (FOVs) from 7 mice. **(C)** Schematic analysis paradigm for detecting synchronous Ca^2+^ activity in meningeal macrophages. **(D)** Synchronous Ca^2+^ events among meningeal macrophages within a FOV. Left: Mean projection of an example FOV showing Ca^2+^ activity in distinct macrophages (colored/numbered). Scale bar 50 μm. Right: Spatial map of macrophages exhibiting synchronous Ca^2+^ activity. Lines connect macrophages with synchronized Ca^2+^ activity, and colors indicate the extent of Ca^2+^ event synchronization. **(E)** Distribution of distances across macrophage pairs showing different numbers of synchronous Ca^2+^ events. **(F)** Linear regression showing poor correlation between distances of macrophage with synchronized Ca^2+^ activity and event delay. **(G)** Proportion of macrophages exhibiting a specific synchronized interaction (cells interacting only with the same subtype: P-P/NP-NP; cells interacting only with a different subtype: P-NP; mixed (cells interacting with the same and different subtypes).

Macrophage intercellular communication, including synchronized activity, potentially across connected macrophage networks, has been implicated in maintaining tissue homeostasis and immune function (*18, 21, 48, 49*). To investigate meningeal macrophage intercellular interactions, we characterized spatiotemporal relationships between Ca^2+^ events in distinct cells within each FOV. We compared temporal factors, including the relative onset latency (t0_A_ – t0_B_) between different macrophages and the duration of the first occurring event (t1_A_ – t0_A_) (**Figure 4C**). We also compared the distances between macrophage pairs exhibiting concurrent events and the number of synchronous events (**Figure 4D**). Finally, we calculated the proportion of perivascular and non-perivascular macrophages exhibiting synchronous Ca^2+^ events. Across all FOVs, 49.3% of macrophages exhibited co-activation over 0–300 μm with minimal distance–delay correlation (**Figure 4E, F**), suggesting that spatial proximity does not influence event synchronicity. Both macrophage subtypes exhibit temporally coincident Ca^2+^ elevations (**Figure 4G**), consistent with a shared synchronization driver. The frequency of synchronous Ca^2+^ events detected could have been influenced by their duration (i.e., the longer the events, the higher the chance of detecting simultaneous event pairs). However, the duration of a given event was a poor predictor of the number of simultaneous events (**Figure 4H**).

### Dural perivascular macrophage Ca^2+^ activity is tuned to behaviorally-driven dural vasomotion

Brain border-associated macrophages in the leptomeninges and related parenchymal perivascular spaces regulate pial arterial vasomotion indirectly by affecting vessel stiffness (*6*). Yet, interaction between vascular-associated macrophages residing in the dura mater, the outermost meningeal layer, and dural vasomotion remains unknown. We therefore imaged the dural vasculature (using a TRITC-Dextran tracer) together with macrophage Ca^2+^ activity and used a Generalized Linear Model (GLM) approach to investigate functional interaction between dural perivascular macrophage Ca^2+^ signals and dural vessel dynamics (**Figure 5A**). Dural arteries constrict during locomotion, while pial arteries dilate (*50*). We analyzed the locomotion-associated responses of 86 meningeal vessels (32 FOVs from 5 mice) and identified a subset (22%; n=19) in which the diameter changes were well fit by a GLM with locomotion state as a predictor. Of these, we identified 74% (n=14) as dural vessels based on their GLM’s negative coefficients consistent with constriction (**Figure 5B and 5C**). Next, we fitted the Ca^2+^ signal observed in perivascular macrophages associated with these dural vessels (n=35) to a GLM using the diameter changes as a predictor variable. Overall, the Ca^2+^ activity of 83% (n=29) of these dural macrophages was well predicted by the model (average deviance explained across all well-fit macrophages: 0.43±0.15, mean±SD). Analysis of the macrophage-vascular models’ beta coefficients revealed two distinct interactions. About half of the macrophages (55%, n=16) exhibited negative coefficients (i.e., increase and decrease in Ca^2+^ activity associated with dural vasoconstriction and recovery, respectively; **Figure 5B and 5D**). The remaining macrophages (45%, n=13) exhibited positive coefficients (i.e., decrease and increase in Ca^2+^ activity in response to dural constriction and recovery, respectively; **Figure 5B and 5E**). The coefficients for increased and decreased macrophage Ca^2+^ activity peaked near zero delay relative to the vasoconstriction and were not statistically different (**Figure 5D-E and 5H**). These data provide evidence that dural perivascular macrophages are functionally coupled to locomotion-driven dural vasomotion, either responding to or mediating it.

**Figure 5.**
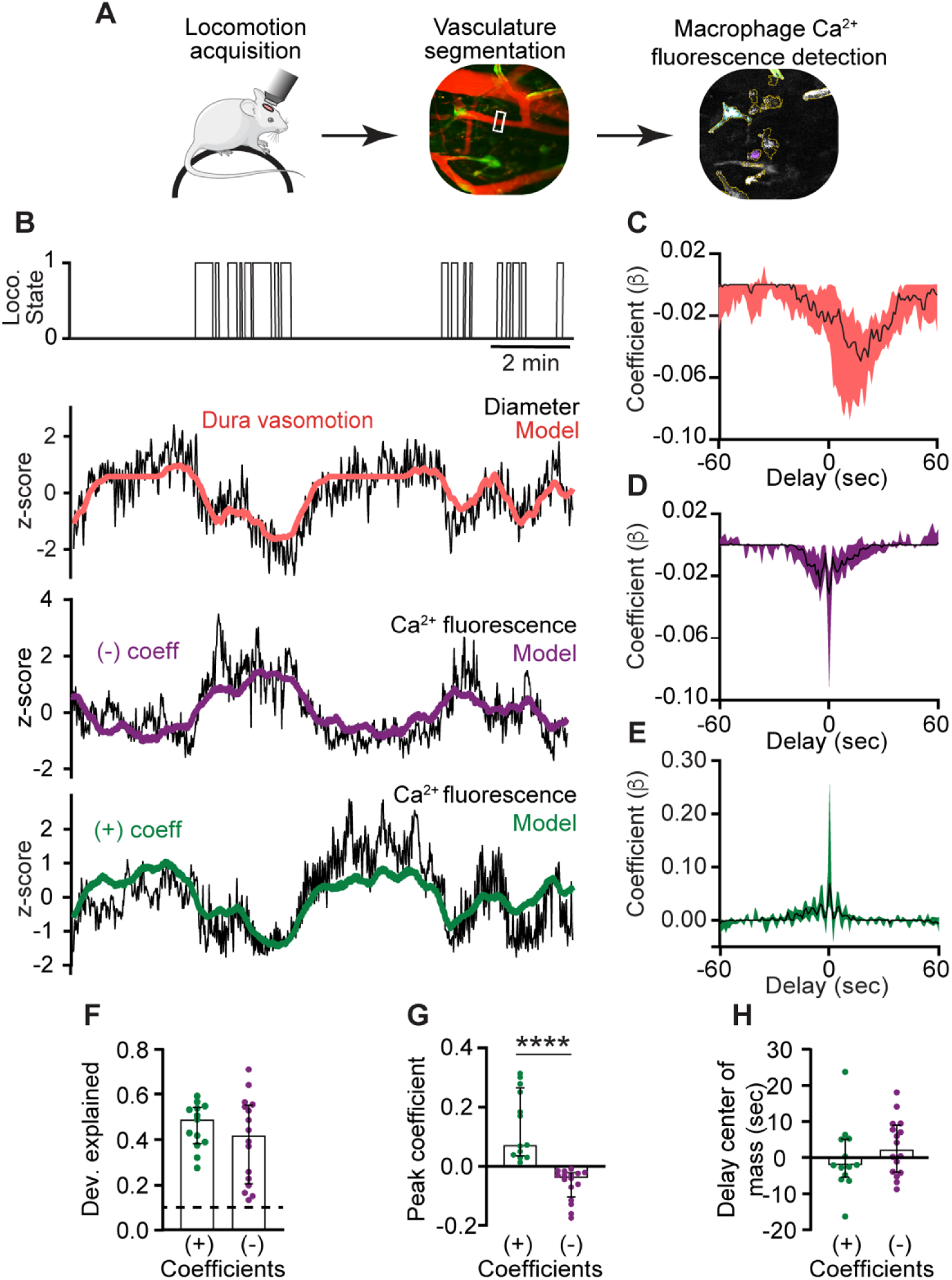
Ca^2+^ signals of dural perivascular macrophages are functionally coupled to behaviorally driven dural vasomotion. **(A)** Experimental paradigm: Locomotion data were acquired during imaging in awake-behaving mice. Behaviorally-evoked changes in meningeal vessel diameter were obtained using segmentation of vessels labeled with a tracer during macrophage Ca^2+^ and further tested for coupling with Ca^2+^ signals. **(B)** Example data of macrophages with Ca^2+^ activity tuned to locomotion-related dural vessel vasomotion. Locomotion bouts (top trace) and dural vessel diameter (black trace) that was well fit by a GLM (red line) using locomotion state as a predictor. Note vasoconstriction during locomotion, indicating a dural vessel. The two bottom traces depict the Ca^2+^ signal (black traces) of distinct meningeal macrophages that were well-fit by a GLM using dural vessel diameter as a predictor, showing either a negative coefficient (purple) or a positive coefficient (green). **(C)** Temporal profile of dura vessels GLM coefficients averaged across all well-fitted vessels (n=19). Traces represent the median across all well-fit ROIs, with shaded regions indicating IQR. **(D, E)** Temporal profiles of GLM coefficient values for macrophages’ Ca^2+^ activity averaged across all well-fitted cells (purple, negative coefficient, n=16; green, positive coefficient, n=13). Traces represent the median across all well-fit macrophages, with shaded regions indicating IQR. **(F)** The deviance explained (goodness-of-fit estimates) of the vessel diameter data included in the GLM used to predict macrophage Ca^2+^ activity were not statistically different for macrophages showing negative (n=16) and positive (n=13) coefficients, suggesting similar interaction levels. **(G)** Comparison of peak positive and negative coefficients of well-fit macrophage Ca^2+^ activity/vasomotion GLMs. **(H)** Center of mass of GLM coefficients indicating that dural vessel diameter changes drive bidirectional changes in fluorescence signal at zero delay. Data (F-H) are median ± IQR. ****p<0.0001 (Mann-Whitney U-test). Data from 32 FOVs from 5 mice.

### An acute aberrant pro-inflammatory brain hyperexcitability event drives diverse Ca^2+^ dynamics in meningeal macrophages

CSD is a slowly propagating depolarization of neurons and astrocytes that drastically disrupts transmembrane gradients and cortical synaptic activity. This aberrant brain hyperexcitability event has been linked to parenchymal inflammation and pain in migraine, traumatic brain injury, and stroke (*26–28*), and could also affect meningeal macrophages (*51*). In anesthetized mice subjected to a single CSD episode, a small subset of meningeal macrophages undergoes morphological changes resembling an inflammatory state (*26*). Given the direct anatomical and functional connections between the brain and meninges (*11, 52*) and the involvement of increased Ca^2+^ influx in macrophage inflammatory activation (*53*), we asked whether CSD drives intracellular Ca^2+^ elevations in meningeal macrophages. We used a pinprick stimulus in the frontal cortex to trigger a single CSD episode in awake mice (*29, 54*) and characterized the related changes in meningeal macrophage Ca^2+^ dynamics. In each experiment, we verified CSD induction based on the associated acute meningeal deformation and/or pial vasoconstriction observed in mice ((*29*) and **Video S2**). We studied CSD-related changes in Ca^2+^ dynamics in 249 macrophages (perivascular, n=64; non-perivascular, n=185; 13 FOVs from 10 mice). For each cell, we compared Ca^2+^ event rates during the passage of the CSD wave (1 min) and the PostCSD period (30 min) to baseline (PreCSD, 30 min) to assess acute and persistent Ca^2+^ responses, respectively. Given the low Ca^2+^ activity observed under steady state and the likelihood that no spontaneous Ca^2+^ elevations occur during the brief period of the CSD event, we considered cells to be either acutely activated or to exhibit an unchanged Ca^2+^ response (i.e., not activated). For studying more prolonged changes during the post-CSD period, we characterized cells as exhibiting persistently increased (event rate > 2× PreCSD), decreased (event rate < 0.5× PreCSD), or unchanged responses. These criteria were used to account for large, observable variations from baseline activity, while also minimizing the influence of spontaneous fluctuations observed in naïve mice. While consistent with previous studies on macrophages in different tissues (*22*), these changes were not intended to represent definitive biological criteria for Ca^2+^ activation and inhibition, but rather a descriptive categorization based on comparable individual cell data.

Using these criteria, we detected both acute and persistent Ca^2+^ activity changes following CSD (**Figure 6 A-E and Video S2**). While smaller subsets of meningeal macrophages exhibited acute (21.3%, n=53) and/or persistent increases (22.1%, n=55) in Ca^2+^ activity, we observed a persistent decrease in the majority of cells (58.6%, n=146). An acute increase was observed more often in peri-vascular macrophages (perivascular, 32.8%, n=21; non-perivascular, 17.3%, n=32, **Figure 6F**). Persistent changes in Ca^2+^ activity were similarly observed in peri- and non-perivascular macrophages (increases; perivascular, 28.2%, n=18; non-perivascular, 18.4%, n=37; decreases; perivascular, 50.0%, n=32; non-perivascular, 61.6%, n=114, **Figure 6G**). The macrophages’ propensity to develop a persistent Ca^2+^ increase was unrelated to their acute response (**Figure 6H**), suggesting that the mechanisms underlying these two temporal responses are distinct. However, cells that showed no acute activation were more likely to exhibit decreased Ca^2+^ activity post-CSD (**Figure 6H**). Finally, we observed that macrophages exhibiting a persistent increase in Ca^2+^ activity had lower baseline activity than those showing a persistent decrease (**Figure 6I**), suggesting that this post-CSD response is influenced by the macrophages’ basal Ca^2+^ activity.

**Figure 6.**
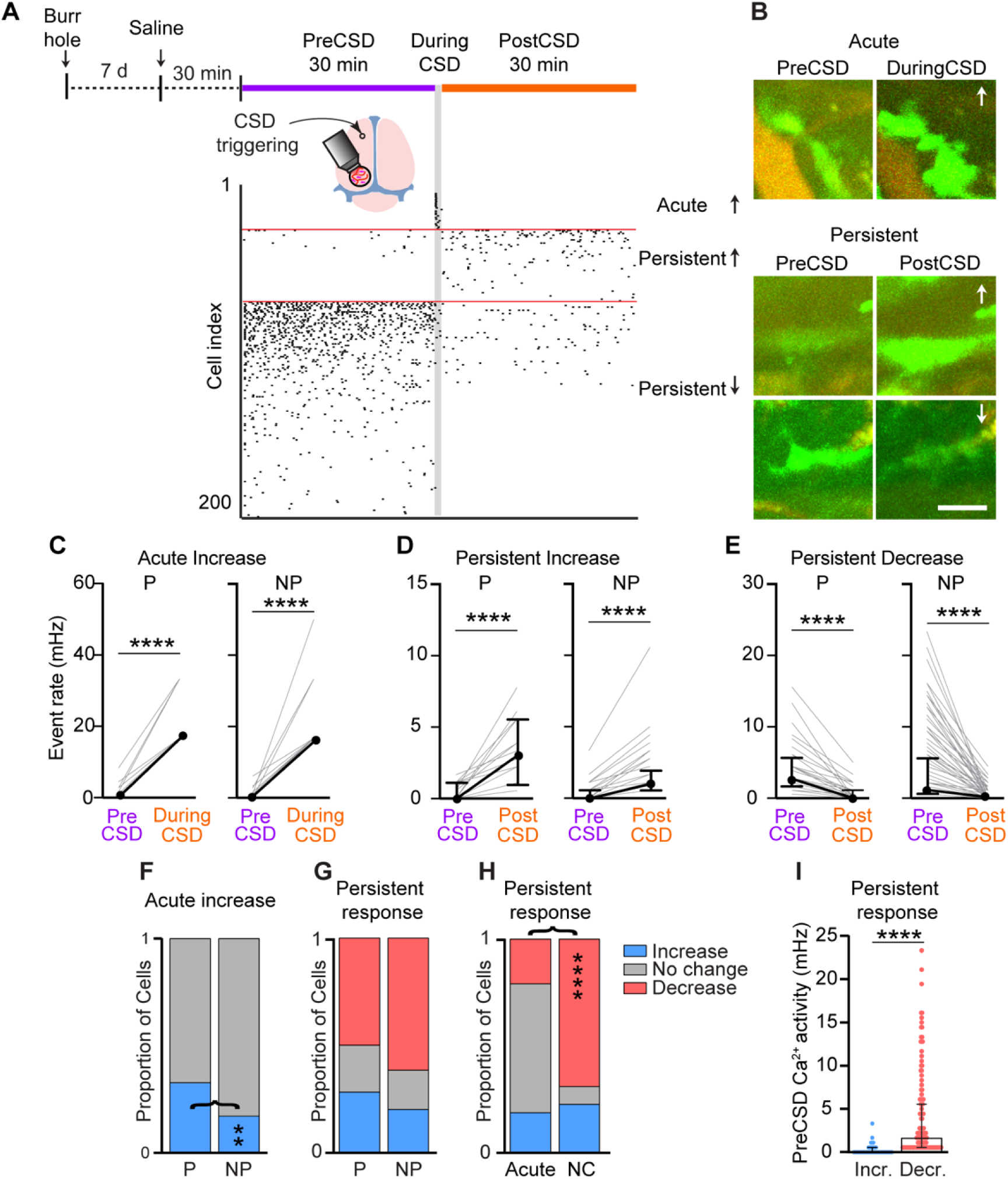
Diverse meningeal macrophage Ca^2+^ dynamics following cortical spreading depression (CSD). **(A)** Experimental setup and example data: (top) A small burr hole was drilled above the frontal cortex 7 days before Ca^2+^ imaging. Mice were pretreated with saline 30 min before imaging baseline macrophage Ca^2+^ activity (30 min, PreCSD). CSD was then induced with a pin prick, and macrophage Ca^2+^ activity was assessed during CSD (1 min during CSD), and post-CSD (30 min, post-CSD). (Bottom) A raster plot of macrophage Ca^2+^ activity showing the acute and persistent increases and persistent decrease in response to CSD. **(B)** Example of macrophage Ca^2+^ fluorescence changes following CSD. Images depict the mean projection over the specific experimental timeline. Arrows indicate an increase or a decrease in Ca^2+^ activity. Scale bar 50 μm. **(C)** Individual responses of perivascular (P, n=21) and non-perivascular (NP, n=32) macrophages showing an acute increase in Ca^2+^ activity. **(D)** Individual response of P (n=18) and NP (n=37) macrophages exhibiting a persistent increase in Ca^2+^ activity. **(E)** Individual responses of P (n=32) and NP (n=114) macrophages showing a persistent decrease in Ca^2+^ activity. **(F)** Proportion of P and NP macrophages showing an acute increase in Ca^2+^ activity or no acute change. **(G)** Proportion of P and NP macrophages showing a persistent increase, decrease, or no change in Ca^2+^ activity. **(H)** Proportion of macrophages displaying distinct persistent responses stratified based on their acute response. **(I)** Baseline (PreCSD) Ca^2+^ activity in macrophages exhibiting persistent increased (n=55) or decreased activity (n=146). Data (C-E) are median ± IQR. ****p<0.0001 (Wilcoxon signed rank test). Data (F-H) represents the proportion of cells, **p<0.01; ****p<0.0001 (Chi-square test). Data (I) are median ± IQR. ****p<0.0001 (Mann-Whitney U-test). CSD data from n=64 perivascular cells and n=185 non-perivascular cells; 13 FOVs from 10 mice.

### CGRP/RAMP1 signaling mediates CSD-evoked persistent increase in meningeal macrophage Ca^2+^ activity

Many meningeal macrophages are localized near peptidergic, CGRP-expressing sensory axons (*16*). In the wake of CSD, cortex-to-meninges signaling enhances the responsiveness of meningeal sensory neurons that could drive CGRP release from their peripheral nerve endings (*29, 54, 55*). CGRP-expressing sensory neurons regulate tissue immunity and meningeal macrophage function via the CGRP/RAMP1 neuroimmune axis (*16, 56*). We therefore asked whether the CSD-related changes in meningeal macrophage Ca^2+^ dynamics we observed involve CGRP/RAMP1 signaling. We pretreated mice with the selective RAMP1 antagonist BIBN4096 and then imaged meningeal macrophage Ca^2+^ activity (42 cells; perivascular, n=14; non-perivascular, n=28; 3 FOVs from 3 mice) before and after CSD. As expected, RAMP1 blockade did not affect CSD triggering (*57*). Compared with the control saline treatment, RAMP1 blockade also did not reduce basal macrophage Ca^2+^ activity (**Figure 7B**). Blocking CGRP/RAMP1 signaling neither affected the incidence of acute increases in Ca^2+^ activity (**Figure 7A and 7C**) nor the magnitude of that response (**Figure 7D**). RAMP1 antagonism, however, inhibited the CSD-evoked persistent increase in the macrophage’s Ca^2+^ activity, without affecting the incidence of the persistent decrease (**Figure 7E**). The data suggest that in the wake of CSD, the CGRP/RAMP1 axis is responsible for the prolonged enhancement of Ca^2+^ signaling in a subset of meningeal macrophages, which could potentially mediate their pro-inflammatory response.

**Figure 7.**
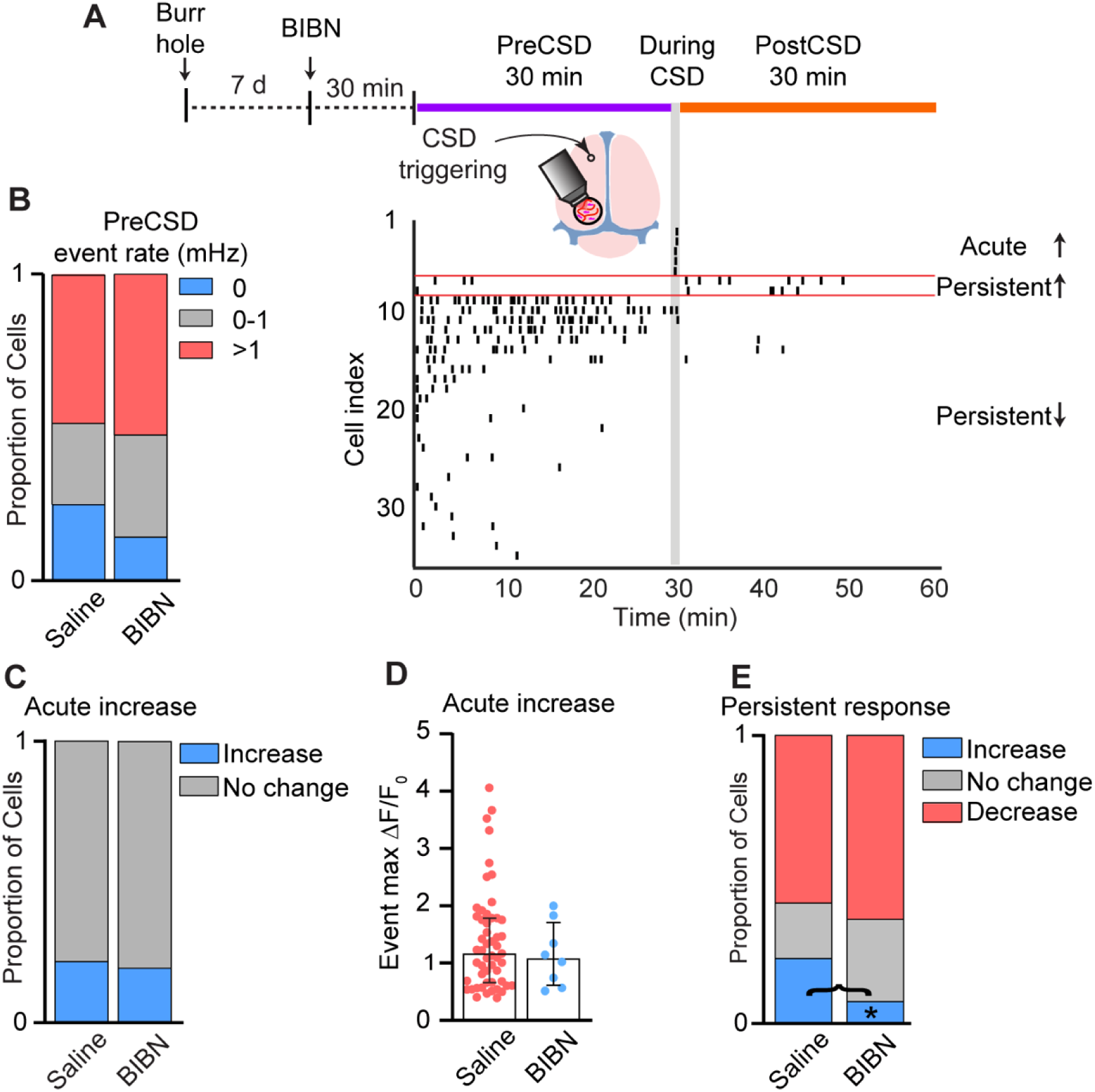
CGRP/RAMP1 signaling mediates CSD-related persistent increase in meningeal macrophage Ca^2+^ activity. **(A)** Experimental setup and example data: (top) A small burr hole was drilled above the frontal cortex 7 days before Ca^2+^ imaging. Mice were pretreated with the RAMP1 antagonist BIBN4096 (BIBN) 30 min before imaging baseline macrophage Ca^2+^ activity (30 min, PreCSD). CSD was then induced with a pin prick, and macrophage Ca^2+^ activity was assessed during CSD (1 min during CSD), and post-CSD (30 min, post-CSD). (Bottom) A raster plot of macrophage Ca^2+^ activity showing the acute and persistent increases and persistent decrease in response to CSD. **(B)** RAMP1 inhibition does not affect the baseline (PreCSD) event rate. Data compared between macrophages imaged in saline-treated mice (n=249 cells, 13 FOVs from 10 mice) and BIBN-treated mice (n=42 cells, 3 FOVs from 3 mice). **(C)** RAMP1 antagonism does not affect the CSD-evoked acute increase in macrophage Ca^2+^. Proportion of macrophages showing an acute response (increase vs. no change) in saline- and BIBN-treated mice. **(D)** RAMP1 antagonism does not affect the magnitude of the acute macrophage Ca^2+^ signal. Event max DF/F_0_ in macrophages showing an acute Ca^2+^ increase in saline-treated mice (n=53 cells) and BIBN-treated mice (n=8 cells). **(E)** RAMP1 antagonism distinctly inhibits the persistent increase in macrophage Ca^2+^ activity post CSD. Proportion of macrophages showing persistent increase, persistent decrease, or no persistent change in saline-treated mice (n=249 cells) and BIBN-treated mice (n=42 cells). **(F)** Data (B, C, E) represents the proportion of cells. *p<0.05, Fisher’s exact test. Data (D) are median ± IQR.

## Discussion

Resident macrophages in the brain meninges are essential for maintaining brain homeostasis, regulating central nervous system immune surveillance, and mediating neuroimmune responses under pathological conditions (*12, 14, 16, 45, 58, 59*). Macrophages rely on Ca^2+^ signaling to mediate many of their functions (*17–23*), yet remarkably little is known about the Ca^2+^ response properties of meningeal macrophages at steady state and disease. Here, using two-photon microscopy in awake behaving Pf4^Cre^:GCaMP6s reporter mice, we provide a foundational landscape of meningeal macrophage Ca^2+^ dynamics. We describe a heterogeneity of meningeal macrophage Ca^2+^ signals at steady state and in response to CSD, an aberrant cortical hyperexcitability event associated with migraine, traumatic brain injury, and stroke. Our data suggest that macrophages in discrete perivascular and interstitial non-perivascular meningeal niches exhibit several distinct Ca^2+^ signal properties at steady state. We further demonstrate Ca^2+^ activity in dural perivascular macrophages, which is tuned to behaviourally driven dural vasomotion. Finally, we describe opposing Ca^2+^ responses of meningeal macrophages following CSD, and demonstrate the contribution of the CGRP/RAMP1 axis in mediating CSD-evoked persistent Ca^2+^ elevations.

The exact link between the distinct Ca^2+^ signal properties of meningeal macrophage subsets observed herein and their homeostatic function remains to be established. The lower event magnitude and noisier signal observed in dural perivascular macrophages may reflect functional interactions with the pulsation dynamics of dural vessels (*50*). Indeed, by combining vascular and macrophage Ca^2+^ imaging, we demonstrate a tight temporal association between the diameter fluctuations of the dural vessels and the Ca^2+^ signal of their associated macrophages in awake, locomoting mice. Direct vascular-macrophage coupling may underlie this interaction, involving macrophages sensing vascular-related mechanical changes via Piezo1 signaling (*60*). Functional vascular-macrophage interaction may also involve mural cells as intermediators (*45, 61*). Studying whether macrophage Ca^2+^ signaling regulates dural vasomotion will require an experimental approach that has yet to be developed, enabling selective manipulation of perivascular dural macrophages. The paucity of ongoing Ca^2+^ activity in perivascular macrophages situated in the leptomeninges we observed supports recent findings that subdural perivascular macrophages indirectly affect pial and parenchymal vasomotion via extracellular matrix remodeling (*6*).

Intracellular Ca^2+^ signal propagation has been observed in various non-excitable cells, such as astrocytes (*62*). We show that the majority of meningeal macrophages, including perivascular and interstitial cells, exhibit intracellular Ca^2+^ signals that propagate throughout the entire cell, suggesting microdomain elevation of intracellular Ca^2+^ following release from internal stores. By characterizing the spatiotemporal relationships between Ca^2+^ signals in distinct cells, we also demonstrate synchronous events that are independent of spatial proximity, suggesting that synchronous Ca^2+^ activity is not driven by intercellular communication. Further studies will be required to resolve the exact source of synchrony. Interestingly, our data indicate that synchronized events involve both peri- and non-perivascular macrophages, despite distinct Ca^2+^ elevation patterns, suggesting that these meningeal macrophage subtypes similarly sense and respond to signals underlying synchronous activity.

Cortex-to-meninges signaling involves a relatively slow flow of soluble molecules within the cerebrospinal fluid that reach the subarachnoid space and then advance via arachnoid cuff exit points into the dura mater (*11, 52*). Our findings of acute Ca^2+^ elevation in a subset of extrasinusoidal perivascular dural macrophages coinciding with the CSD event suggest a rapid transfer of soluble signaling factors released from a hyperexcitable cortex across all meningeal layers (*28*). Nevertheless, we cannot exclude a mechanically driven macrophage response to the acute meningeal deformation produced by the neuronal and glial swelling and shrinkage of the cortical extracellular space during CSD (*29, 63, 64*). Our data also indicate a delayed, prolonged increase in Ca^2+^ signaling in a relatively small subset of macrophages post-CSD, which could underlie their proinflammatory-like morphological change (*51, 53*). Our findings also support the view that meningeal neuroimmune CGRP/RAMP1 axis serves as a mechanism responsible for this macrophage Ca^2+^ response, potentially via the activation of macrophage RAMP1/CLR receptor complex (*16*) by CGRP released from sensitized meningeal afferent axons (*29, 54, 65*). Whether the relatively small subset of meningeal macrophages featuring increased Ca^2+^ signaling serves a protective role (*66, 67*) or a proinflammatory, destructive function (*68*) remains to be elucidated. Intriguingly, our data points to a persistent decrease in macrophage Ca^2+^ activity post-CSD, not involving CGRP/RAMP1 signaling, as the most prevalent response. Further studies are needed to determine whether this reduction in Ca^2+^ activity reflects altered viability or reduced immune function that could interfere with the macrophage’s ability to restore homeostasis and dampen local inflammation (*69*).

There are several limitations to our study. First, while PF4^Cre^-based labeling has been shown to target brain border-associated macrophages, we cannot fully exclude the possibility that in a small subset of meningeal dendritic cells, monocytes, and T cells that have low-level PF4 expression (10), GCaMP6 was also expressed, leading to a Ca²⁺ signal. Nonetheless, a recent study using PF4^Cre^mTmG mice failed to detect EGFP reporter expression above background in any other meningeal cells by flow cytometry (*70*). Second, to enhance Ca^2+^ event detection, we downsampled the movies to ∼1 Hz.

We therefore could have missed fast Ca²⁺ transients or microdomain activity. Third, in our study, we imaged Ca^2+^ activity in extrasinusoidal meningeal macrophages. It is therefore possible that these cells exhibit distinct response properties compared to the subset of dural macrophages associated with the dural sinuses (*8*). Finally, our study used a pharmacological approach to determine whether CGRP/RAMP1 receptor signaling mediates macrophage Ca^2+^ responses to CSD. We acknowledge that this approach does not allow us to establish a specific role for macrophage CGRP signaling, given the possibility that CGRP/RAMP1 signaling in other meningeal vascular or immune cells (*10, 16, 71*) may indirectly affect the macrophage Ca^2+^ response.

## Conclusions

We provide a detailed characterization of macrophage Ca^2+^ dynamics in homeostatic meninges, thereby expanding our understanding of their biological diversity. The coupling of dural perivascular macrophage Ca^2+^ signals and dural vasomotion may represent a unique homeostatic functional dural macrophage-vascular unit that controls dural perfusion. The diversity of meningeal macrophage Ca^2+^ responses to CSD further highlights the complexity of brain-to-meninges neuroimmune signaling and meningeal macrophage function in neurological disorders such as migraine, traumatic brain injury, and stroke. Our study also provides essential genetic and data analysis tools to further understand the molecular signaling underlying macrophage function at steady state and neuropathological conditions.

## Methods

### Animals

All experimental procedures were approved by the Beth Israel Deaconess Medical Center Institutional Animal Care and Use Committee. Experiments were conducted on adult Pf4^Cre/wt^:TIGRE2.0^GCaMP6s/wt^ Ca^2+^ reporter mice (8-17 weeks) (9 males, 5 females). Mice were generated by crossing Pf4^Cre^ (C57BL/6-Tg (Pf4-icre) Q3Rsko/J, Jackson laboratory, Strain #008535) mice with GCaMP6s^fl/fl^ (Ai162D; B6.Cg-Igs7^tm162.1(tetO-GCaMP6s,CAG-tTA2)Hze^/J, Jackson laboratory, Strain #031562) mice. Animals were genotyped by Transnetyx Inc.

### Surgical procedures

Animals were anesthetized using isoflurane in 100% O_2_ (induction: 3%; maintenance: 1.5-2%) and placed on a heating pad with a rectal probe attached to a stereotaxic frame to monitor animal body temperature during surgery. Animals received dexamethasone (8 mg/kg, i.p.) and Meloxicam SR (4 mg/kg, s.c.) to reduce inflammation and improve surgical outcomes. An eye ointment was used to prevent ocular drying. Mice were implanted with a titanium headpost and a 3 mm glass cranial window (1.5 mm lateral and 2 mm posterior to Bregma) over an intact dura covering the left posterior neocortex (*29*). Immediately after surgery, the mouse cage was placed on a water-circulating heating pad for faster recovery. Animals were then single-housed with access to a running wheel and a hut and allowed to recover for at least one week.

### Wheel running acclimation

After the cranial window surgery, mice were allowed to recover for at least a week. To reduce stress associated with head-fixation during imaging and habituate to wheel running, the mice received multiple training sessions (10 min to 1 h over 3–4 days). In each session, the mouse was placed on a 3D printed running wheel, with its headpost attached to two clamps, and allowed to locomote freely.

### Two-photon Ca^2+^ imaging

Awake-behaving mice were head-fixed to the running wheel by its headpost (**Figure 1**). We used a two-photon microscope (Neurolabware) with a Nikon 16X, 0.8 N.A. objective to acquire images at 15.5 Hz with digital zoom set at 4X (312 x 212 µm^2^ FOV). A MaiTai laser set to 920 nm with 25–40 mW power was used to excite fluorescence. The Scanbox package for MATLAB (Neurolabware) was used to control the microscope and acquire images and wheel running data. To image the meningeal vasculature, mice were administered 70 kDa TRITC-Dextran tracer (50 mg/kg, i.v.; Sigma-Aldrich).

### Behavioral tracking during two-photon imaging

We recorded running speed in MATLAB using a custom-made encoder (Arduino) coupled to the 3D-printed running wheel. See below for details of analyses of behavioral variables.

### Induction of CSD and pharmacological treatment

For CSD induction, a 1 mm burr hole was drilled at the frontal bone (1.5 mm anterior to the cranial window) to allow access to the brain cortical surface. A small amount of silicone elastomer (Kwik-Cast, WPI) was placed to cover the burr hole opening, and the animal was left to recover for at least one week. CSD was induced using a brief 2-second cortical pinprick (*29*). CSD induction was confirmed by the identification of a short-lasting meningeal deformation and/or transient pial constriction (*29*). On the experimental day, 30 minutes before baseline recording, mice were pretreated with the selective CGRP/RAMP1 antagonist BIBN4096 (0.3 ml, 1 mg/kg, i.p., Tocris) (*55*) or 0.3 ml of saline (Vehicle control).

### Quantification and statistical analysis

#### Two-photon imaging movie processing

We used a discrete Fourier transform to perform rigid registration to correct for translation changes caused by brain motion during locomotion. Movies were then downsampled to 1.03 Hz. Locomotion signals were detected as described (*43*). All image processing and locomotion signal extraction were performed in MATLAB 2021b (Mathworks).

#### Ca^2+^ signal detection pipeline

For detecting macrophage Ca^2+^ signals, we used the Activity Quantification and Analysis (AQuA2) platform that implements an event-based approach with advanced machine learning techniques for temporal and spatial segmentation of Ca^2+^ fluorescence events ^38^. Importantly, this computational platform captures event dynamics beyond traditional ROI-based approaches. The following user-defined parameters were input: 0.49 µm/pixel spatial resolution, 1.03 Hz temporal resolution, 1-second minimal event duration detection; window of event size between 25% (125 µm^2^/ 523 pixels) and 150% (750 µm^2^/ 3140 pixels). Every event and cell identified was followed by a manual visual check. Subsequently, each cell was labelled as perivascular or non-perivascular according to its location relative to vessels. Morphological features (area, perimeter, circularity) and spatiotemporal aspects of the Ca^2+^ signals (i.e., ΔF/F_0_ dynamics, frequency, amplitude, duration) were used to analyze cell-specific characteristics and Ca^2+^ activity profiles. We employed the AQuA2 automatically-generated function (*44*) to characterize intracellular Ca^2+^ propagation. Intercellular Ca^2+^ activity was evaluated by analyzing temporally co-occurring events (synchronized event pairs), as well as their corresponding spatial localization in the FOV. All post-processing of AQuA2-generated data was performed using MATLAB 2021b (Mathworks).

#### Clustering of Ca^2+^ dynamics

We used a Savitzky-Golay filter to detrend and smooth Ca^2+^ activity traces. Polynomial order and frame size were optimized for each cell by selecting the combination of parameters that yielded a better signal-to-noise ratio. From the filtered signals, we extracted the dominant frequency using Fast Fourier Transform, and peak counts using minimum peak prominence (threshold of 10% signal amplitude range). The optimal number of clusters was determined using the Elbow method, after which k-means clustering was applied to group cells based on their signal characteristics.

#### Analysis of CSD-related changes in macrophage Ca^2+^ dynamics

For analyzing the effects of CSD on macrophages’ Ca^2+^ dynamics, we divided the activity of each cell into three phases: ‘PreCSD’ (minutes 0 – 30), ‘DuringCSD’ (minutes 30 – 31), and ‘PostCSD’ (minutes 31 – 61). Ca^2+^ event rates during CSD and PostCSD were compared with those of the PreCSD baseline to evaluate acute and persistent Ca^2+^ responses, respectively. Ca^2+^ responses were categorized as increased (event rate > 2x PreCSD), decreased (event rate < 0.5x PreCSD), or unchanged.

#### Analysis of locomotion

We extracted running speed (cm/sec) from the wheel encoder. To infer the locomotion state, we first concatenated all velocity signals obtained from a given mouse across all experiments and trained a two-state Hidden Markov Model using the MATLAB function ‘hmmtrain’. Then, the locomotion state was inferred for each individual imaging run by applying the MATLAB function ‘hmmviterbi’ with the model trained on the concatenated data. Locomotion bouts were defined as periods when the locomotion state was sustained for at least two seconds.

#### Vascular signals

We calculated changes in vascular diameter by first generating a maximum-intensity projection of the red channel and then drawing polygons (ROIs) around each vessel. Subsequently, for each frame, pixels inside each ROI were extracted, and a Radon transform was applied to get a 1D vessel profile. Radon transform of the first frame was used as a reference to normalize the results of subsequent frames, resulting in a time series of normalized diameter traces.

#### General linear models (GLM)

We investigate the functional interaction between dural perivascular macrophage Ca^2+^ activity and dural vasomotion by fitting a Gaussian GLMs using the GLMnet package (MATLAB 2021b), with elastic net regularization (α=0.01) and ten-fold cross-validation. We employed a two-step modeling. The first model was used to identify dural vessels by evaluating the correlation between changes in vessel diameter and locomotion state, and classifying dural or pial vessels based on their vasoconstriction and vasodilation dynamics, respectively (*50*). The second modeling step evaluated the relative contribution of perivascular dural Ca^2+^ activity signals to changes in dural vessel diameter. All signals were downsampled to 1.03 Hz to match the frame rate used when extracting fluorescence Ca^2+^ signals in AQuA2. To allow potential anticipatory or delayed responses of diameter-to-locomotion or Ca^2+^ fluorescence-to-diameter, the time interval used to analyze the response to the predictor was set from -60 s to +60 s. The GLM was trained on 75% of the data, and all predictions and model performance reported are from the remaining 25% testing set. A threshold of 0.1 goodness-of-fit deviance explained was set. For each predictor temporal shift, a response coefficient was generated. The centroid delay between the predictor and response was calculated, weighted by the absolute value of the coefficient (i.e., center-of-mass, COM).

#### Statistical analysis

All statistical analysis was performed using GraphPad Prism 10.4 and MATLAB 2021b. Data were analyzed using a Wilcoxon matched-pairs signed rank sum test or a Mann-Whitney U-test. Distribution of categorical data was analyzed using the Chi-square or Fisher’s exact test. P-values are indicated as follows: *p<0.05, **p<0.01, ***p<0.001, ****p<0.0001.

## Data availability

All data needed to evaluate the conclusions in the paper are present in the paper.

The code used to analyze the data in this study was deposited on the Levy Lab GitHub account. Code for movie processing is available at: https://github.com/levylabheadache/MovieProcessing/tree/SCN;

Code for locomotion analysis is available at: https://github.com/levylabheadache/Locomotion/tree/SCN;

Code for post Aqua2-processing is available at: https://github.com/levylabheadache/Aqua2Processing;

Code for vascular segmentation is available at: https://github.com/levylabheadache/Vasculature/tree/SCN;

Code for GLM is available at: https://github.com/levylabheadache/GeneralLinearModel_Macrophages

Any additional information is available from the corresponding author upon request.

## Abbreviations

AQuA2: Activity Quantification and Analysis
Ca^2+^: Cytoplasmic calcium
CGRP: Calcitonin gene-related peptide
CSD: Cortical spreading depolarization (depression)
CX3CR1: CX3C Motif chemokine receptor 1
GLM: General Linear Model
NP: Non-perivascular
P: Perivascular
Pf4: Platelet factor 4
RAMP1: Receptor activity modifying protein 1
TRITC: Tetramethylrhodamine isothiocyanate

## Authors information

D.L. conceived the project. S.C-N and D.L. wrote the manuscript. S.C-N performed two-photon imaging and data analysis with help from D.L. S.C-N, C.W., and A.G. performed surgeries.

## Supplementary video legends

**Video S1. Macrophage Ca^2+^ dynamics in homeostatic meninges of awake mice**

Example two-photon imaging showing Ca^2+^ activity in perivascular and non-perivascular meningeal macrophages.

**Video S2. Acute and persistent changes in meningeal macrophage Ca^2+^ activity in response to CSD.**

Example two-photon imaging of meningeal macrophage Ca^2+^ activity at baseline, during, and following CSD. The arrow indicates a macrophage showing an acute Ca^2+^ elevation, the arrowhead depicts a delayed and persistent Ca^2+^ elevation, and the asterisk, a macrophage showing a persistent decreased Ca^2+^ activity.

